# The genetic architecture of the pepper metabolome provides insights into the regulation of capsianoside biosynthesis

**DOI:** 10.1101/2023.09.27.559835

**Authors:** Julia Nauen, Pasquale Tripodi, Regina Wendenburg, Ivanka Tringovska, Amol N. Nakar, Veneta Stoeva, Gancho Pasev, Annabella Klemmer, Velichka Todorova, Mustafa Bulut, Yury Tikunov, Arnaud Bovy, Tsanko Gechev, Dimitrina Kostova, Alisdair R. Fernie, Saleh Alseekh

## Abstract

*Capsicum* (pepper) is among the most economically important species worldwide, the fruit accumulates specialized metabolites with essential roles in plant environmental interaction and potential health benefits. However, the underlying genetic basis of their biosynthesis remains largely unknown. In this study, we developed and assessed both wild genetic variance and a bespoke mapping population to determine the genetic architecture of the pepper metabolome. The genetic analysis provided over 30 metabolic quantitative trait loci (mQTL) for over 1100 metabolites. We identified 92 candidate genes involved in various mQTL. Among the identified loci, we described and validated by transient overexpression a domestication gene cluster of eleven UDP-glycosyltransferases involved in monomeric capsianoside biosynthesis. We additionally constructed the biosynthetic reactions and annotated the genes involved in capsianoside biosynthesis in pepper. Given that differential glycosylation of acyclic diterpenoid glycosides contributes to plant resistance and acts as anticancer agents in humans, our data provide new insight, and resources for better understanding the biosynthesis of beneficial natural compounds to improve human health.

## Introduction

Pepper (*Capsicum* spp.) is one of the most important cultivated crops in the Solanaceae family with a global gross production value of $37.6 billion in 2021 (fao.org/faostat/en/#compare). Pepper fruit has a broad range of pharmacological applications and is consumed worldwide as a vegetable, fruit, and spice (Srinivasan, 2005; Chen et al., 2009; Meghvansi et al., 2010). *Capsicum* comprises more than 36 species; however, *C. annuum*, *C. chinense*, *C. frutescens*, *C. pubescens*, and *C. baccatum* species have been domesticated over the course of many centuries, and are predominantly cultivated across different pepper-producing regions (Walsh and Hoot, 2001; Carrizo García et al., 2016; Liu et al., 2023). Each species has a distinctive phytochemical and morphological diversity (Antonio et al., 2018; Naves et al., 2022; Liu et al., 2023). The most common species, *C. annuum*, is said to have originated from Mexico and migrated to Africa and Asia via Mesoamerica, and reached Europe ultimately (Hayano-Kanashiro et al., 2016; Tripodi et al., 2021).

Plant-specialized metabolites are critical for biotic and abiotic interactions with the environment and are crucial for human nutrition and health, with numerous medical applications (Martin and Li, 2017; Martin et al., 2011; Jiang et al., 2021). Varieties of *C. annuum* vary in pungency due to the presence or absence of capsaicinoids, unique compounds that produce a burning sensation due to somatosensory and viscerosensory perception (Frias and Merighi, 2016; Heiser and Smith, 1953; Nelson, 1919). Capsaicinoids serve as protectants against plant diseases and are known to promote cell death of malignant cells in humans (Sánchez et al., 2006; Batiha et al., 2020). Next to essential vitamins, minerals, and nutrients, peppers are also rich in a number of phytochemicals such as carotenoids, capsaicinoids, flavonoids, ascorbic acid, and tocopherols (Marín et al., 2004; Kozukue et al., 2005). These compounds are all known to prevent inflammatory diseases associated with oxidative damage to maintain optimum health (Martin et al., 2011; Martin and Li, 2017; Jiang et al., 2021). Compounds exclusively produced in the genus *Capsicum* are capsaicinoid’s sweet analogs capsinoids (Sutoh et al., 2006), capsidiols (Stoessl et al., 1972), capsanthin (Davies et al., 1970), capsicosides (Yahara et al., 1994), and the acyclic diterpenoid glycosides known as capsianosides (Izumitani et al., 1990). Capsianosides have a wide range of glycosylation and hydroxylation states (Fig. S1) and can exist as monomers or dimeric esters (Izumitani et al., 1990; Iorizzi et al., 2001; De Marino et al., 2006; Lee et al., 2008, 2006, 2009). As a result, capsianosides have been shown to act as antioxidants with anticancer properties in humans (Chilczuk et al., 2020) and to contribute to defense-related responses in plants due to their Ca^2+^ chelating properties (Macel et al., 2019; Bacon et al., 2017; Hashimoto et al., 1997). Although a wide range of secondary metabolites including the capsianosides have been reported in pepper (Wahyuni et al., 2014, 2013, 2011; Naves et al., 2022; Wu et al., 2023), the genetic basis underlying their biosynthesis pathways is poorly exploited. Combining metabolomics with association and linkage mapping has revealed that it is possible to decipher the genetic architecture of a variety of crop plants, including maize, rice, and wheat, as well as non-cereal species such as lettuce and tomato (Alseekh et al., 2020a, 2015; Zhu et al., 2018; Zhang et al., 2020a; Zhan et al., 2020; Wen et al., 2014). This increasingly important research field facilitates phenotype screening without the construction of transgenic plants (Fernie and Schauer, 2009).

In this study, we used two mapping populations, a genome-wide association study (GWAS) panel alongside a biparental backcross inbred line (BIL) population, and explored the fruit metabolic diversity in multiple experiments (Fig. S2). Metabolite GWAS (mGWAS), and metabolite quantitative trait locus (mQTL) mapping revealed genetic architectures responsible for the natural variations in the composition of secondary metabolites in the pepper populations. We identified 32 QTL and 92 novel genes associated with the abundance of 20 different compound classes. To demonstrate the power of our approach and resources provided here, we focused on a fruit-specific metabolic hotspot on chromosome 9 associated with monomeric capsianoside, combining the QTL analysis from both GWAS and biparental BIL population. We identified a gene cluster of 11 UDP-glycosyltransferases (UGTs) involved in metabolic diversity of the monomeric capsianosides in pepper. The function of the UGTs were verified by transient overexpression in pepper and tobacco. Furthermore, we discovered evidence of selection of these UGT genes during pepper domestication by analyzing the genetic architecture of the hotspot QTL. Finally, given the importance of these metabolites both in terms of plant disease resistance and the nutritional importance of these compounds in the human diet, these findings may contribute to the development of elite pepper varieties which possess increased disease resistance and enhanced nutritional value.

## Results

### Developing and genomic composition of the biparental BIL and GWAS populations

The biparental BIL population was developed by crossing *C. chinense* cv. CGN22793 (donor parent) with *C. annuum* cv. Marconi giallo (recurrent parent) to generate 117 BILs each carrying several introgressions from *C. chinense* (Fig. S3). The genotyping by sequencing (GBS) analysis of the BIL resource yielded 10,914 single nucleotide polymorphisms (SNP) with 648 to 2865 introgressions per chromosome (Fig. S4, S5). In the BIL population, there were 157.8 (median: 147) introgressions with an average length of 0.186 Mbp (median: 0.072 Mbp) per introgressed segment (Figure S4c,d). Further, analysis of the genomic composition revealed an average of 7.8 % and 87.4 % of donor and recurrent parents’ genomes, respectively, with less than 3 % heterozygosity (2.2 % missing data; Fig. S5).

In parallel to the BIL population, we explored the genetic diversity of 162 *C. annuum* accessions collected from 62 diverse locations across six Balkan countries. Figure 1 shows the genetic diversity and population structure analysis of the GWAS panel. GBS discovered 77,401 SNPs across 12 chromosomes, providing genome-wide coverage (Fig. 1a,b,d). Further, the data showed three subpopulations, which could be primarily attributed either to their pungency or the kapia and pumpkin fruit shape (Fig. 1b,c).

**Figure 1.**
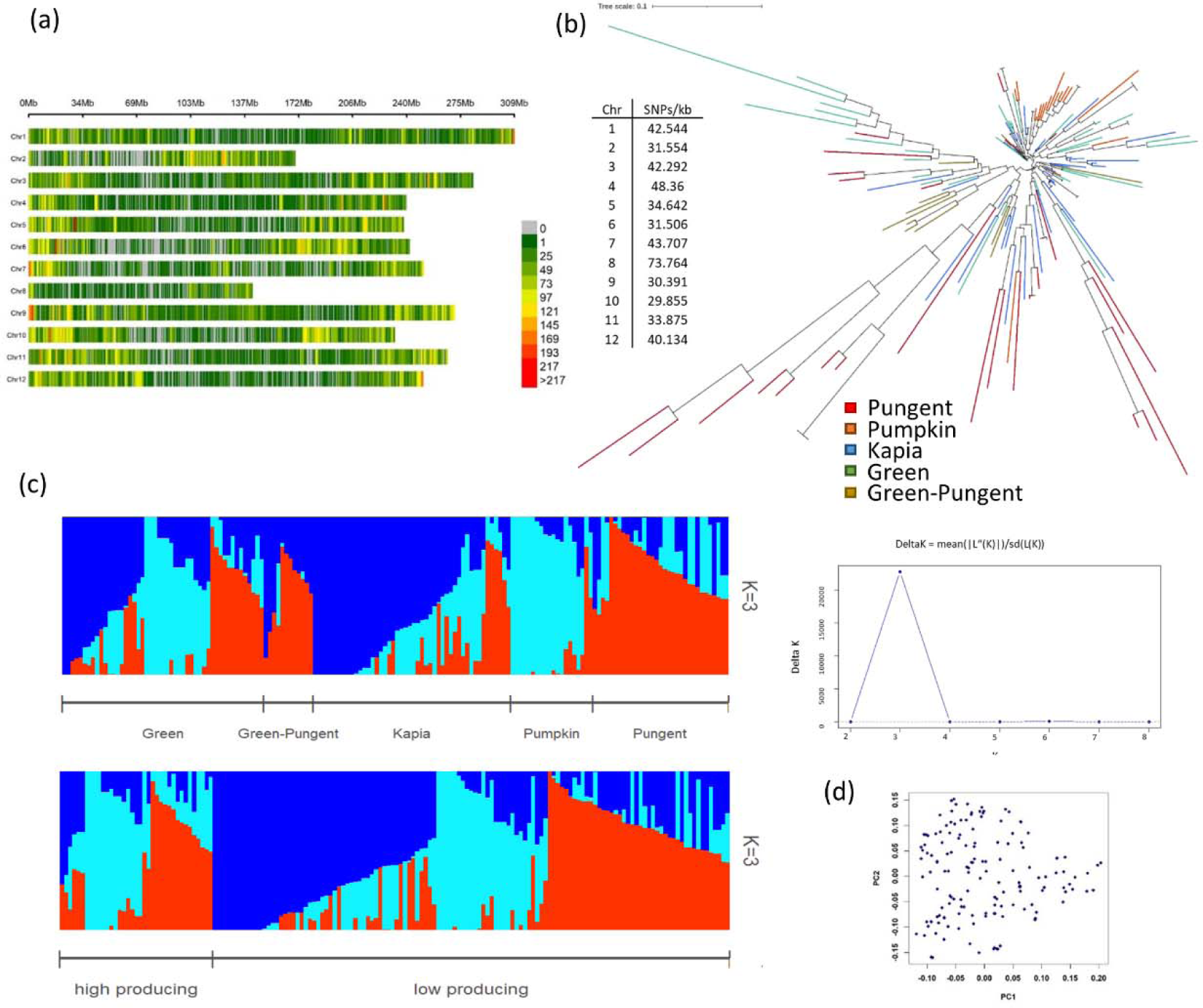
Genetic architecture of the genome-wide association study (GWAS) population. a, SNP density within one megabase window size generated by genotyping-by-sequencing. b, Phylogenetic analysis of 162 *C. annuum* accessions using 77,401 SNPs. c, Hierarchical population structure analysis of 162 accessions of *C. annuum* from the Balkans. Using 77,401 single nucleotide polymorphisms, the number of subpopulations was most likely inferred at K = 3. Population structure at K = 3 based on Q matrix assigning accessions either to primary morphological groups or production of Capsianoside IX. Colours represent subpopulation. d, PCA of GBS data from GWAS panel.

### Metabolic diversity in the GWAS and BILs populations

In order to explore the chemical diversity in pepper fruit, we grew both the GWAS and BIL populations for multiple experiments, collected fruit materials, and subject them to ultra-high performance liquid chromatography coupled to mass spectrometry (UHPLC-MS) for metabolite profiling (see materials and methods). Using UHPLC-MS we were able to detect and quantify about 1100 metabolites across both populations including 456 flavonoids, 422 acyclic diterpenoids, 153 other phenolics, and 59 other compounds including lipids and fatty acids (Data S1 – S5, Fig. S6).

To estimate the genetic:environmental variation of the observed metabolites, we calculated the broad-sense heritability (H^2^) for all metabolic traits, in the GWAS population H^2^ was > 50 % for 88 % of the compounds, of which 9.5 % of the metabolites had H^2^ > 90 % (Data S1, S2, S5). The relative difference in the abundance of 13.6 % of the compounds had a variation > 100-fold. Analyzing the chromatograms in an untargeted manner, 5362 mass features were identified with 87.2 % of the mass features discerned variation > 50 % whereas 11.1 % of the mass features showed more than 90 % variation, and the rest 2.5 % of the mass features showed > 100-fold variation. Amongst the compounds which showed high H^2^ and high metabolic variation were acyclic diterpenoid derivatives (e.g., capsianoside V, III, X-2, IX, IV, II + malonyl), flavonoids (e.g., apigenin-6-*C*-pentoside-8-*C*-hexoside, luteolin-6,8-di-*C*-hexoside, quercetin-3-*O*-trisaccharide), capsaicinoids (e.g., nonivamide, capsaicin) and sesquiterpenoid (e.g., Cadina-1(6),4-diene). Similar results were also obtained from the BIL populations under two growing conditions, the glasshouse (experiment 1) and polytunnel (experiment 2; Data S4, S5). Of these metabolites 73 % had H^2^ > 50 %, with 6.1% compounds > 80 % and 23 % higher 100-fold variation. Analyzing H^2^ of 2067 common compounds identified under both growing conditions in an untargeted manner, 74.8 % of the compounds had H^2^ > 50 % with 6.8 % of compounds > 80 % H^2^. The variation among the common metabolites was for 24.2 % of the identified compounds higher than 100-fold.

To investigate the metabolic diversity further, principal component analysis (PCA) was performed for all metabolic traits (Fig. S6). In the case of data from the GWAS panel, PC1 and PC2 explained 39.8 % and 6.5 % of the trait variance, respectively (Fig. S6b). Figures S6c and d show the metabolic variation of the BILs from the experiment one and two. In this case, PC1 could explain 28.7 % and 29.7 % of the variation, while PC2 could explain 13.3 % and 13.9 % of the variation in both experiments, respectively (Fig. S6c,d). PC1 distinguished the parental lines while PC2 distinguished lines with unique metabolic variation.

### Genetic basis of the pepper fruit metabolome

Having estimated the fraction of variation that is genetically determined, and revealed a high number of measured metabolites that can potentially be mapped to mQTL, we next performed mGWAS for the GWAS panel and mQTL mapping for the BILs to reveal the genetic basis of pepper fruit metabolome (GWAS SNPs Data S6, BIL SNPs Data S7). We used mixed linear modeling (MLM), and R-QTL to detect significant associations in GWAS and the linkage mapping populations, respectively. In total, we identified 1941 lead SNPs for 156 annotated metabolites and 13,792 lead SNPs for 1278 non-annotated compounds in the GWAS panel (Data S8). Linkage mapping in the BIL population identified 551 SNPs with LOD score > 15 for 265 annotated metabolites, and 4764 significant SNPs for 2376 non-annotated compounds. Pleiotropic analysis which marks overlapping associations in a 50 kb window identified 45 candidate genes for several overlapping significant loci in both BIL and GWAS populations (Fig. 2, Data S9).

**Figure 2.**
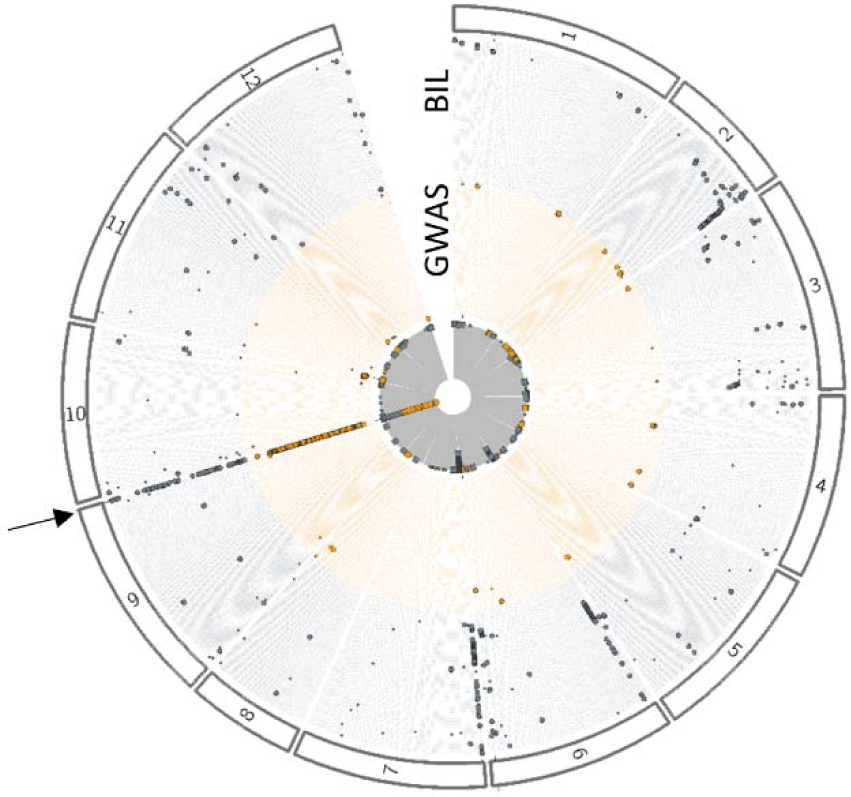
Pleiotropy analysis of GWAS and BIL panel. Genome-wide association study (GWAS; orange; 173 semi-polar compounds) and linkage mapping (blue; 190 semi-polar compounds) performed of fruit pericarp samples. The pleiotropic map identifies regions of the genome that have significant single nucleotide polymorphisms (SNPs). Each dot denotes a significant GWAS or linkage mapping hit that has been mapped to a specific genomic region. The inner grey circle shows the total number of significant SNPs mapped to the genomic area, the outer ring represents the number of chromosomes. The UGT cluster (arrow) is mapped by a total of 132 and 84 semi-polar compounds from the GWAS and BIL populations, respectively.

The highest sum of associations was observed at the bottom of chromosome 9 (Fig. 2, arrow).

In order to demonstrate the power of both GWAS and linkage mapping populations to reveal candidate genes controlling metabolic traits, we first mapped the capsaicinoid content (metabolites that contribute to pepper fruit pungency) of both populations (Fig. S7). Both the GWAS and QTL mapping panels revealed significant associations for capsaicinoids (e.g., dihydrocapsaicin, capsaicin, and nonivamide; Fig. S7). One of the QTL regions spanning 205.2 kb (149.95 Mb – 150.15 Mb, LOD = 15.15) on chromosome two contained 13 genes, including the acyltransferase 3 *AT3/PUN1*. This gene has previously been described as the causal enzyme catalyzing the final condensation reaction of the biosynthesis of capsaicin (Qin et al., 2014). The same QTL was additionally validated by mapping the fruit pungency scores (see materials and methods) in the GWAS panel (Figure S7b). This result demonstrated the power of both populations to identify key genes involved in pepper metabolism. As a further example, we have identified a new locus on chromosome two 1.6 Mb downstream controlling the abundance of capsaicinoids. This QTL region overlapped in both populations and harbored 3 genes (151.68 Mb – 151.86 Mb, LOD = 16.39) including *CA02g20190* coding for an MYB transcription factor that could be used for further validation and follow-up studies (Data S9, S11). Furthermore, Wu *et al.,* (2023) recently validated the transcription factor *CaMYB12-like* (Wu et al., 2023), a QTL found in our mapping populations (Fig. S8), in association with epicatechin, quercetin, naringenin, and its glucoside, and capsianoside I (Data S9, S11).

Next, we mined candidate genes that had not previously been identified. For example, pleiotropic analysis identified six QTL, associated with various metabolic traits (Data S9). For instance, on chromosome 1, we identified a UDP-galactose/UDP-glucose transporter at position 56.74 Mb (*CA01g19370*, 56.61 – 56.79 Mb, LOD = 14.6) and a Progesterone 5-beta-reductase at position 57.37 Mb (*CA01g19140*, 57.32 – 57.372 Mb, LOD = 13.74) associated with several flavonoid derivatives such as naringenin-*O*-hexose 2, apigenin 6,8-di-*C*-hexoside, luteolin 6-*C*-hexoside, and quercetin-*3-O*-rutinoside-*7-O*-glucoside. On chromosome 6 at position 234.76 Mb a MYB transcription factor (*MYB36*, *CA00g82220*, 234.59 – 234.81 Mb, LOD = 11.89), a Galactosyltransferase 8 at position 234.85 (*CA06g26600*, 234.81 – 234.96 Mb, LOD = 11.89) and two carbonic anhydrases nectarin-3 at position 234.98 – 234.99 Mb (*CA00g91080, CA00g91090*, 234.96 – 235.06 Mb, LOD = 13.6) were found to associate to naringenin-*C*-hexoside-pentoside, luteolin 6-*C*-hexoside-8-*C*-pentoside, capsianoside V, naringenin-*O*-hexose 2. On chromosome 11, we identified a UDP flavonoid 3-*O*-glucosyltransferase at position 14.2 Mb (*CA11g03000*, 14.18 – 14.4 Mb, LOD = 48.04) associated with naringenin-*O*-hexose among other unannotated compounds (Data S9). Capsianoside III and phenolics (apigenin derivatives, naringenin-O-hexose, caffeic acid 3-glucose, luteolin 6,8-di-C-hexoside) of the BIL population mapped to three cytochrome P450 enzymes on chromosome 12, *CA12g22250* and *CA12g22240* at position 250.376 to 250.379 Mb (LOD = 42.96, 250.375 – 250.383 Mb), and *CA12g22000* 295.8 kb downstream at position 250.67 Mb (LOD = 11.01, 250.579 – 250.699 Mb).

Furthermore, 29 QTL were manually identified: On chromosome 8 at 114.48 Mb, a tocopherol cyclase (*CA08g05970*) was associated with γ-tocopherol (415.19 m /z, 7.88 min) in the GWAS panel (Table S9). Caffeic acid 3-glucoside was linked to a phenylalanine ammonia-lyase (*CA05g20790*) on chromosome 5 at 233.22 Mb. Dimyristoylcapsanthin (1021.823 m/z, 15.1 min) was identified in the BIL population and mapped to a phytoene synthase 1 (*CA04g04080*) on chromosome 4 at 9.8 Mb, next to luteolin 6,8-di-*C*-hexoside and an acyclic diterpenoid derivative and ten phenolics and two capsianoside. A capsanthin/capsorbin synthase (*CA06g22860*) was identified by an association of 3 acyclic diterpenoids and quercetin 3,7-dirhamnoside, quercetin-trihexoside in the BIL population. Eight unannotated compounds next to an acyclic diterpenoid derivative and a sinapoyl derivative mapped to a region on chromosome 2 containing a cluster of 23 1-aminocyclopropane-1-carboxylate oxidase 1 at position 133.7 to 134.3 Mb.

### Pathway construction for capsianoside biosynthesis pathway

Capsianosides are a diverse class of acyclic diterpene glycosides, water-soluble constituents present in pepper. While the chemical identity, anti-herbivore effect, and dietary health benefits have been reported (Macel et al., 2019; Chilczuk et al., 2020), the genetic basis underlying their biosynthesis is yet to be fully comprehended (Fig. S2). In our study, we detected over 400 capsianoside derivatives (Data S5) and were able to identify eight genetic loci controlling their abundance using either association or linkage QTL mapping approaches (Fig. 3, Data S9).

**Figure 3.**
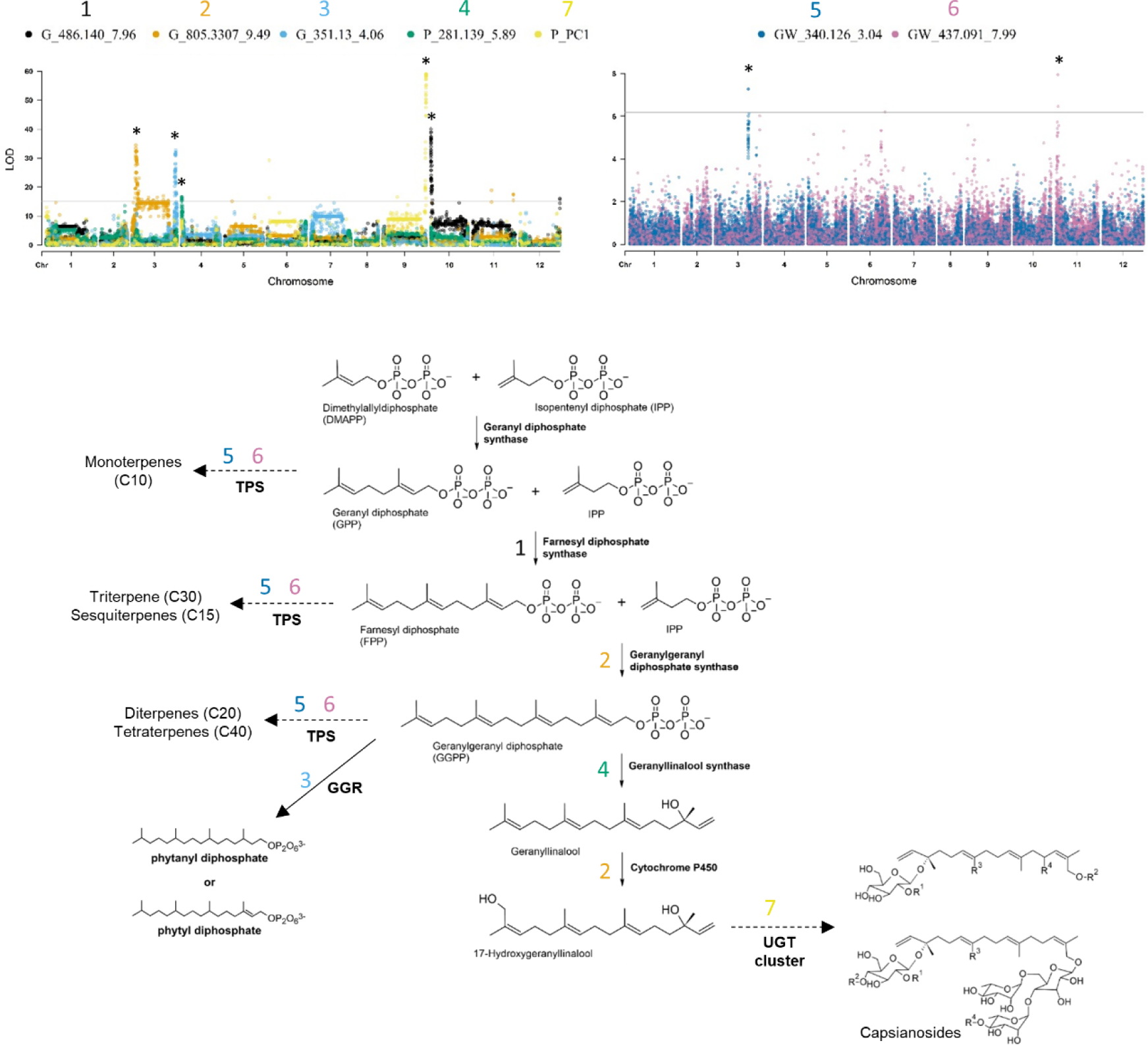
Candidate genes of the capsianoside pathway. Overlayed LOD scores plot of five compounds (1 – 486.140 m/z 7.96 min, 2 – 351.13 m/z 4.06 min, 3 – 281.139 m/z 5.89 min, 4 – 805.331 m/z 9.49 min) and 7 – principal component (PC) 1 and an overlayed manhattan plot of two compounds (5 – 340.126 m/z 3.04 min, 6 – 437.091 m/z 7.99 min) representing the seven quantitative trait loci (QTL) discovered through linkage and association mapping refining the capsianoside pathway: 1 (black) – Farnesyl diphosphate synthase (*CA10g01050*), 2 (orange) – Geranylgeranyl diphosphate synthase (*CA03g17670*) and Geranylgeranyl diphosphate reductase (GGR, *CA03g29990*), 3 (light blue) – Geranyllinalool synthase (*CA04g02600*), 4 (green) – Cytochrome P450 Geraniol 8-hydroxylase (*CA03g17470*), 5 (dark blue) – Terpenoid synthase/cyclase (TPS, *CA03g15870*), 6 (pink) – TPS (*CA11g03240), and* 7 (yellow) – UDP-glycosyltransferase (UGT) cluster. Asterisks indicate QTL of color-coded metabolic features (G = BIL glasshouse Exp. 1, P = BIL polytunnel Exp. 2, GW = GWAS panel).

For example, a 15.1 Mb introgression on chromosome 10 (LOD = 30.88) revealed a farnesyl pyrophosphate synthase (*CA10g01050*) at 817.7 kb among three genes controlling several capsianoside derivatives (Fig. 3, QTL 1). The second QTL in Figure 3 spans 504 kb on chromosome 3 and comprises 41 genes (LOD = 31.33), including geranylgeranyl pyrophosphate synthase (*CA03g17670*) at 25.1 Mb and geraniol-8 hydroxylase (*CA03g17470*) at 24.4 Mb. Another QTL on chromosome 3 (LOD 31.33) spans 144.6 Mb and contains 13 genes, including a geranylgeranyl diphosphate reductase (*CA03g29990*) at 271.89 Mb which is implicated in capsianosides biosynthesis. In addition, QTL four on Figure 3 reveals a geranyllinalool synthase (*CA04g02600*) on chromosome 4 at 636.6 kb. With a LOD of 11.48, the introgression spans 237.3 kb and 22 genes. QTL 5 (-log10(*p*) = 7.28) and 6 (-log10(*p*) = 7.28), both containing terpenoid synthases (TPS), were only found in the GWAS panel. The TPS at position 206.3 Mb on chromosome 3 (*CA03g15870*) is 206.3 kb upstream of the lead SNP, whereas its homolog on chromosome 11 (*CA11g03240*) at position 12.1 Mb is 106.7 kb upstream of the lead SNP. Finally, for QTL 7 (Fig. 3) the PC1 of experiment 2 of the BIL population was used for association mapping (compare Figure 4e, S10), and could resemble the hotspot region on chromosome 9 (Fig. 2), and mapped to an UDP-glycosyltransferase (UGT) gene cluster. In total 23 capsianoside derivatives and 10 flavonoids (BIL exp. 1), 19 capsianoside derivatives, two capsaicinoid derivatives and 7 flavonoids (BIL exp. 2) and in case of the GWAS panel 108 capsianoside derivatives and 14 flavonoids mapped to this region containing 11 UDP-glycosyltransferases.

**Figure 4.**
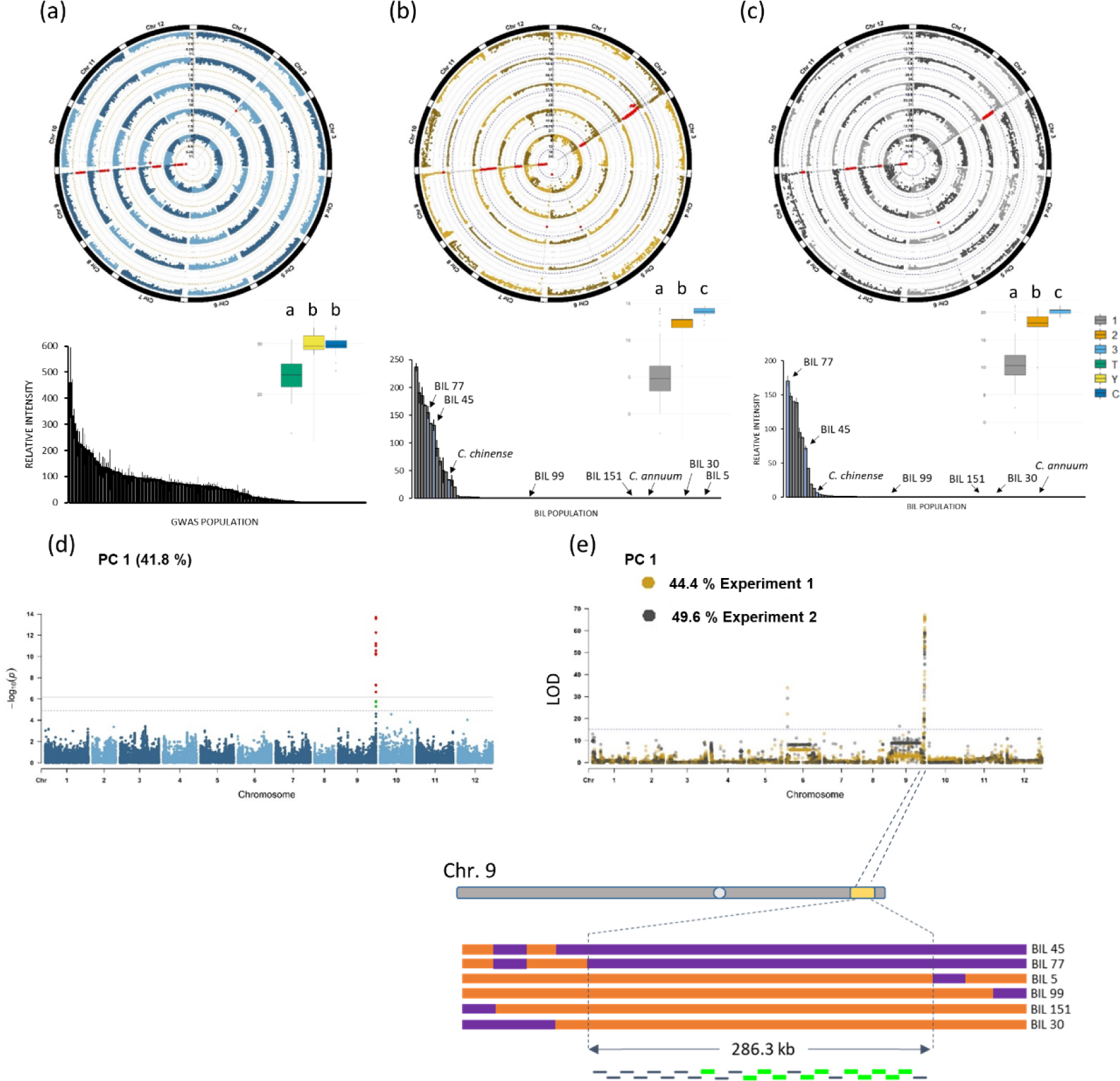
Association of capsianosides to the bottom of chromosome 9. a – c, Circular manhattan plots of genome-wide association study (GWAS; a) and linkage mapping of BILs experiment 1 (b) and 2 (c) of capsianosides (from inside to outside: trihydroxy-phytatetaenoic acid-O-glucose II, dihydroxy-phytatetaenoic acid diglucose, capsianoside X-2 (8.8 min), capsianoside X-2 (10.8 min), capsianoside IX from inside to outside) with boxplots (in log2, 1 = recurrent genome, 2 = heterozygous, 3 = donor genome) and bar plots (mean ± SE) of relative intensity of capsianoside X-2 of 162 accessions and 102 (b) and 100 (c) BILs. Letters indicate significant difference p < 0.01 (student’s t-test). d, e, Mapping of principal component 1 (PC1) of GWAS panel (d) and BIL population (e) experiment 1 and 2. e, A region of 286.3 kilobase pairs on chromosome 9 could be narrowed down to be causal for differential abundance (orange = recurrent genome, purple = donor genome).

### Identification of a gene cluster responsible for capsianoside decoration

The above mQTL analysis identified genetic loci and candidate genes involved in the upstream capsianoside biosynthesis pathway. Intriguingly, the GWAS analysis identified a hot spot mQTL at the bottom of chromosome 9 that influenced a wide range of capsianosides derivatives, specifically the glycosylated derivatives of diterpene backbones (Fig. 4a).

To cross-validate our findings, we also checked the QTL presence in the BIL population (Fig. 4b,c,e). The QTL mapping in the BIL population indeed confirmed the hotspot in two independent experiments (Fig. 4b,c). Further, to investigate the specificity of the mQTL to capsianoside derivatives in this study, we performed PCA analysis using capsianoside derivatives in both GWAS and BIL populations (Figs. 4d,e, S9, S10). Then, PC1 to three scores were used to perform the GWAS and QTL mapping (Figs. 4, S9, S10, Data S12). In GWAS, PC1 explained 41.8 % of capsianosides variance (Fig. S9b). A similar observation was obtained for the BIL population, with 44.4 % and 49.6 % variation in capsianoside explained by PC1 in experiments one and two, respectively (Fig. S10). PCA analysis further indicated that > 85 % of the variation is explained by genetic variance. Interestingly, in both the GWAS panel and BIL population the main peak (-log_10_(*p*) of 13.4 and LOD 67.1 and 59) mapped to the bottom of chromosome 9. Further, using disequilibrium analysis of GWAS (Fig. 5c) and recombination breakpoints in the BIL population (Fig. 4), we were able to narrow down the QTL-hotspot region to 286.3 kb (270.1 – 270.29 Mb), which contains 25 genes (Data S10), including a cluster of 11 UDP-glycosyltransferase (UGT) genes that pinpoint the most likely causative genes.

**Figure 5.**
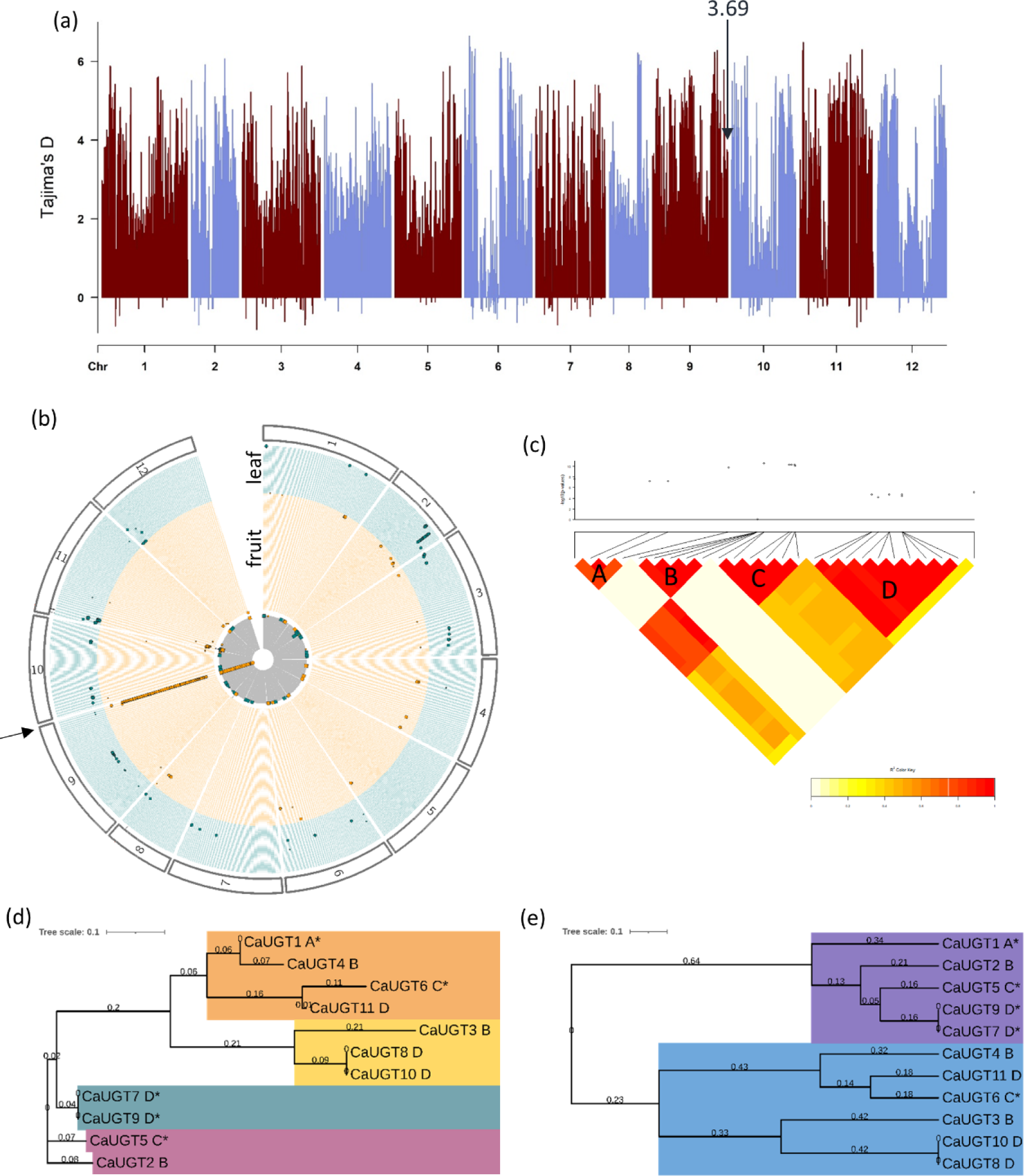
Eleven UDP-glycosyltransferases causal for differential glycosylation of capsianosides in pepper fruit pericarp. a, Tajima’s D was calculated in a 1 Mb non-overlapping bins using 77,401 SNPs from the GWAS panel. The arrow indicates the bin start at position 270 Mb on chromosome 9 where the UGT cluster is located with 114 SNPs resulting in a Tajima’s D of 3.69 indicating a historical selective sweep region. b, Pleiotropic analysis of GWAS of fruit pericarp (orange) and leaves (70 semi-polar compounds; cyan; for further description see Figure 2). c, Genetic linkage analysis as a pairwise linkage disequilibrium heatmap in R^2^ in a ± 50 kb window around candidate genes. d, The cladogram comparing nucleotide sequences (compare Figure S17, S18) and e, amino acid sequences of the UGT cluster. Letters indicate the chromosomal location (linkage blocks see (a)) of the genes, and asterisks indicate genes cloned for transient overexpression.

However, it should be noted that 41 disease-resistance genes are located upstream of the UGT cluster from position 267.7 to 269.4 Mb, and 33 disease-resistance genes are located within or downstream of the UGT cluster from position 270.1 to 271.1 Mb. In total, 25 genes were assigned to one orthogroup (OG0000008, Data S10, S11) and are annotated as disease resistance protein BS2 and are associated with black spot disease in pepper, which is caused by *Xanthomonas campestris* pv. *vesicatoria* (Hibberd et al., 1987).

In addition, to the major hotspot on chromosome 9 controlling a large number of capsianoside derivatives in pepper fruit, the QTL analysis identified additional significant loci specific to the BIL populations (Data S10). The QTL mapped to the bottom of chromosome 2 harboring an introgressed segment from *C. chinense* of 106.4 kb at position 165.14 – 165.28 Mb (LOD = 73.5). This region contains seven genes two of which are predicted to be CRS1-like group II intron splicing factors (*CA02g23580*, *CA02g29200*) orthologous to the tomato *Solyc02g087250* and are promising for further investigation of transgressive segregation among the BIL population. Also, in this region *CA02g29190* encodes a protein of unknown function, whereas the orthologue in *S. lycopersicum* (*Solyc02g087260*), is predicted to be a TMV response-related protein, implying a defense-related role. However, the causality between these genes and the observed phenotype still needs to be investigated.

### Selective sweeps and functional validation of a UDP-glycosyltransferase cluster associated with capsianoside variation in pepper fruit

We concentrated our efforts on the UGT cluster at the bottom of chromosome 9 because it had the greatest number of associations. To confirm that this region was selected by breeders during domestication, we performed a Tajima’s D analysis in 1 Mb non-overlapping bins (Fig. 5a, Data S13). The analysis yielded a Tajima’s D of 3.69 by using 114 SNPs spanning the region from 270 Mb to the telomers, indicating the occurrence of purifying selection. Next, we measured the levels of 102 compounds (33 annotated) in leaves of 156 pepper accessions and performed mGWAS on leaf material by UHPLC-MS to determine whether the mQTL was specific to fruits or also detected in leaves (Fig. 5b, Data S14). None of the capsianoside derivatives measured in leaf tissue were able to be mapped to the bottom of chromosome 9 (Fig. 5b).

The genetic linkage analysis revealed four linkage blocks (Fig. 5c), UGT1 is linked to block “A”, UGT2, UGT3, and UGT4 are linked to “B”, UGT5, UGT6 are linked to “C”, and UGT7, UGT8, UGT9, UGT10, and UGT11 linked to block “D” (Fig. 5c,d). Furthermore, the linkage disequilibrium (LD) analysis indicated that all of the presented linkage blocks (except B) are inherited in LD. Multiple sequence alignment analysis of the UGTs revealed four and two subclusters using the nucleotide and amino acid sequences, respectively (Fig. 5d,e). Interestingly, genes with high sequence similarity are spread across different linkage blocks indicating duplication events.

To test the hypothesis that the UGT gene cluster controls the glycosylation steps in the capsianoside biosynthesis pathway, we cloned the coding sequence of four UGT genes. Independent transient overexpression was performed in *Nicotiana benthamiana*, *C. annuum* (CM334), and *Aarabidopsis thaliana* (Figs. 6, S11 – S13), in order to investigate the role of UGTs in the capsianoside biosynthesis. Four of the UGTs were successfully cloned from *C. chinense* and three from *C. annuum* and localization was determined by a GFP epitope tag driven by CaMV 35S. UGT1 and 6 were found to be cytosolic, while UGT5 and 7/9 were found in the cytosol as well as the chloroplast (Fig. S11, S12).

**Figure 6.**
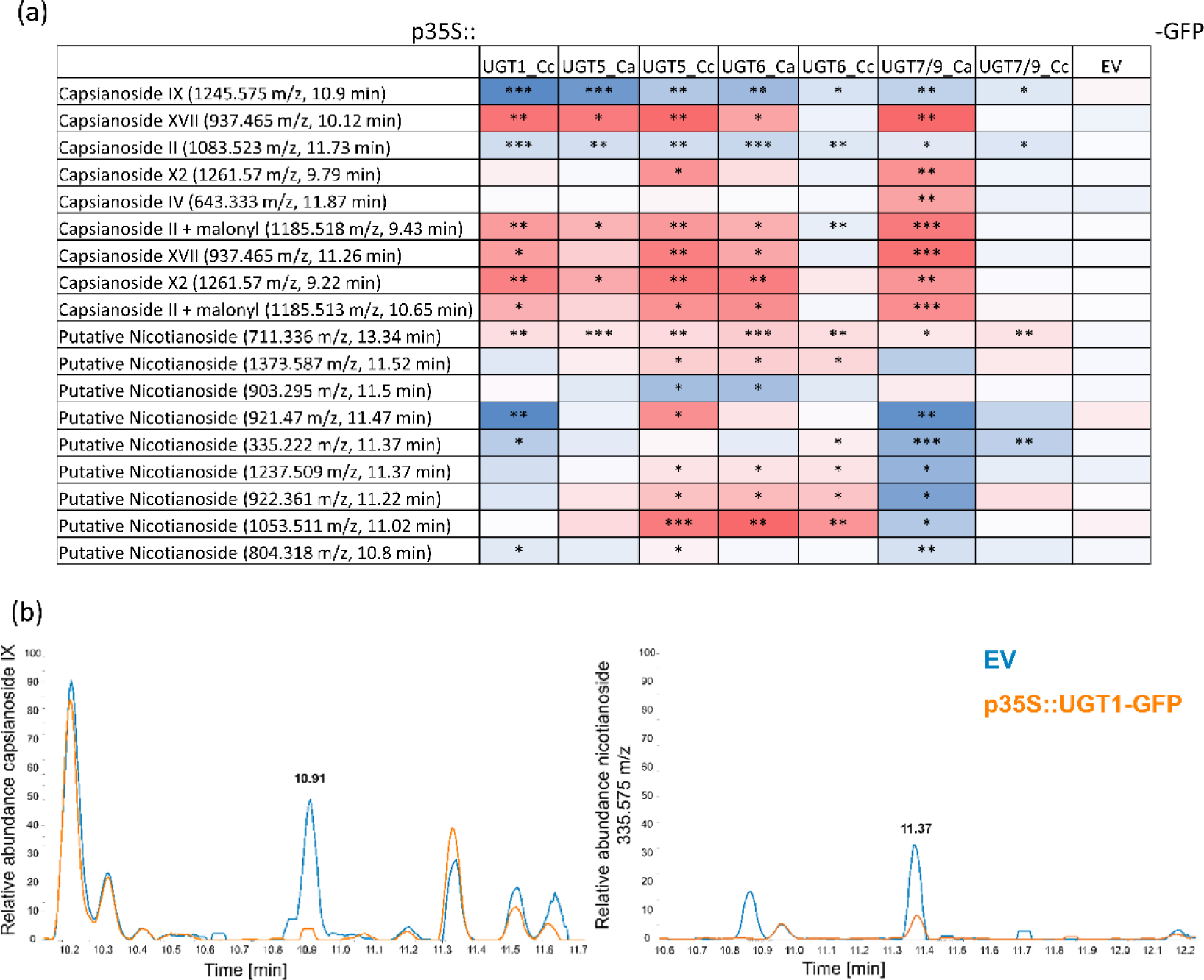
Transient overexpression of leaves of *C. annuum* cv. CM334 and *N. benthamiana*. UGT1, 5, 6, and 7/9 of alleles from *C. annuum* cv. Marconi giallo (Ca) and *C. chinense* cv. CGN22793 (Cc) were cloned (CaMV p35S, C-terminal eGFP) and leaves transformed. a, Heatmap indicates different capsianosides and structurally related nicotianosides. Significances between fold change intensities to empty vector (EV) are indicated by unpaired two-sided Student’s t-test *p < 0.05, **p < 0.01, ***p < 0.001. b, Extracted ion chromatogram of capsianoside IX and a putative nicotianoside comparing EV to p35S::UGT1-GFP from *C. chinense*.

We cloned the coding sequences of four UGT genes to test the hypothesis that the UGT gene cluster controls the glycosylation steps in the capsianoside biosynthesis pathway. To investigate the role of UGTs in capsianoside biosynthesis, independent transient overexpression was performed in *N. benthamiana*, *C. annuum* (CM334), and *A. thaliana* (Figs. 6, S11 - S13). UGT 5, UGT6, and UGT 7/9 (7/9 with 100% sequence identity) were successfully cloned from both *C. chinense* and *C. annuum*, whereas UGT1 was only cloned from *C. chinense*. The localization was determined using a GFP epitope tag driven by CaMV 35S and revealed that UGT1 and 6 are cytosolic, whereas UGT5 and 7/9 are both cytosolic and chloroplastic (Fig. S11, S12). Metabolic profiling of transformed leaves revealed significant differences in glycosylated capsianoside content between transformed pepper leaves over-expressing UGT genes and empty vector (Figs. 6, S14). A similar result was obtained in *N. benthamiana* leaves over-expressing the UGT genes, with especially noticeable changes in nicotianoside derivative content (Diterpene glycosides in *Nicotiana*, Fig. 6). These findings suggest that UGTs are responsible for the decoration of capsianoside derivatives in pepper.

Phylogenetic analysis of multiple UGTs from *A. thaliana* (At), *N. benthamiana* (Nb), and *N. attenuata*, as well as the crop species *Vitis vinifera*, *Punica granatum* (Pg), *Camellia sinensis*, *Glycine max* (Gm), *Sorghum bicolor*, and *Oryza sativa* (Knoch et al., 2018; Wilson and Tian, 2019; Heiling et al., 2021; Dudley et al., 2022) was performed (Fig. 7). CaUGT 1, 2, 5, 9, and 7 cluster with PgUGT722A1 and NbUGT93S1 which belong to group O UGTs. Interestingly, UGT709 belonging to the same group was found to be involved in monoterpene metabolism (Miettinen et al., 2014). CaUGT8, 10, 3, 6, 11, and 4 showed similarity to the outgroup (OG) stated in Wilson and Tian, 2019 (Fig. S15). Next to *A. thaliana*’s UGT80A2 are UGTs from the Poaceae *Brachypodium distachyon*, the Bryophytes *Physcomitrella patens*, *Selaginella moellendorffii*, and the algae *Chlamydomonas reinhardtii*. AtUGT80A2 was previously reported to be involved in the glycosylation of steryl glycosides and acyl steryl glycosides (Schrick et al., 2012; Debolt et al., 2009). However, phylogenetic analysis using 109 UGTs from *N. attenuata* revealed both clusters similar to group O UGTs (Fig. S16). In addition, both clusters of *Capsicum* UGTs are phylogenetically close to di- and triglycoside forming glycosyl transferases and group A UGTs (Fig. 7). *Veronica persica* VpUGT94F1 and GmUGT91H4 were previously shown to be involved in flavonoid and triterpene *O*- and *C*-glycosylation (Ono et al., 2010; Jung et al., 2014; Shibuya et al., 2010; Casas et al., 2016; Itkin et al., 2016).

**Figure 7.**
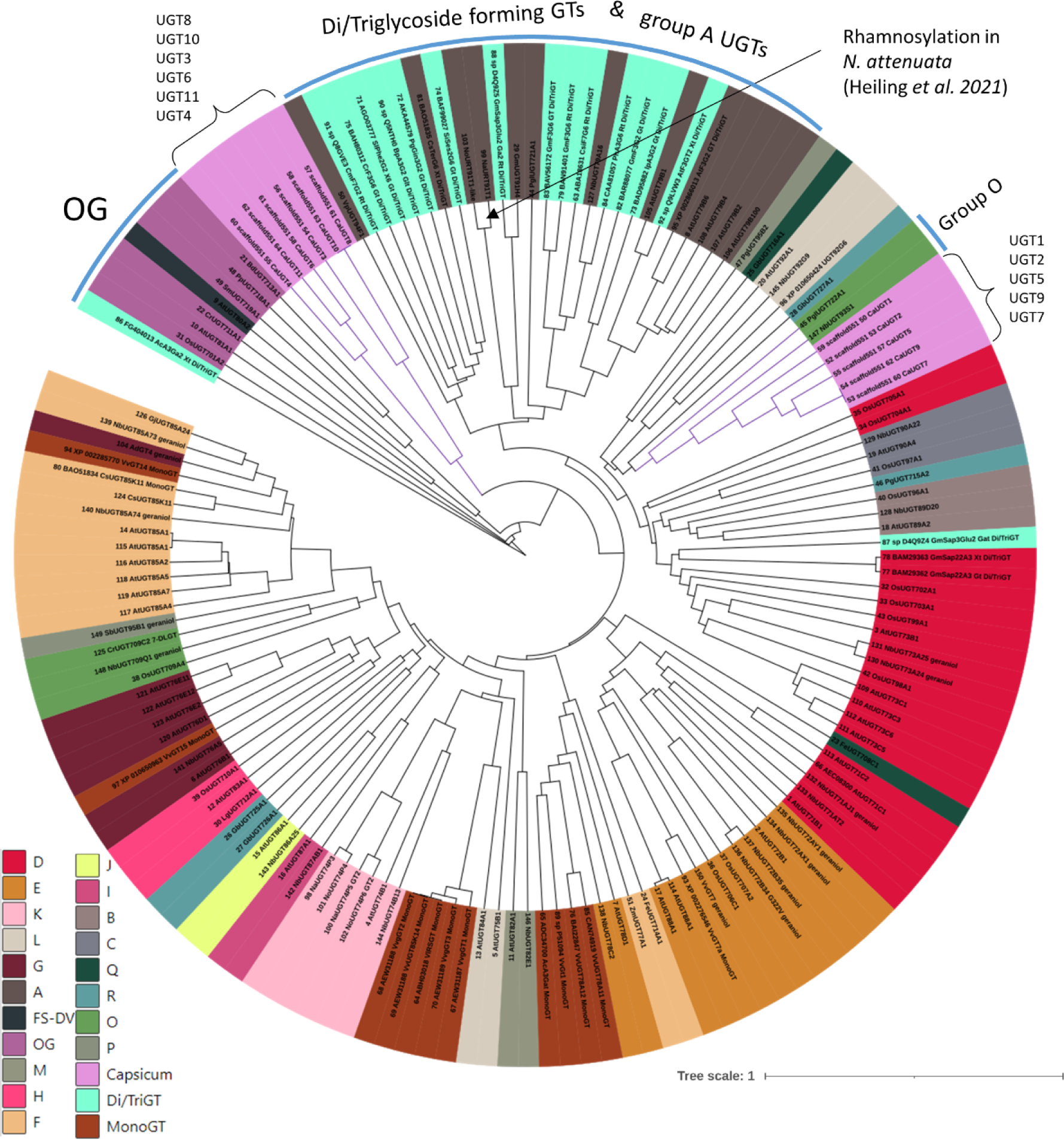
UDP-glycosyltransferase (UGT) 1, 2, 5, 7, 9 cluster next to group O UGTs. Phylogenetic tree based on amino acid sequences of 139 UGTs across different plant species in addition to the eleven CaUGT. *Arabidopsis thaliana* (At), *Nicotiana benthamiana* (Nb), and *N. attenuata* (Na), as well as the crop species Vitis vinifera (Vv), Punica granatum (Pg), Camellia sinensis (Cs), Glycine max (Gm), Sorghum bicolor (Sb), and Oryza sativa (Os). Multiple sequence analysis was performed using Clustal omega (https://www.ebi.ac.uk/Tools/msa/clustalo/) and tree built with interactive Tree of Life (iTOL, https://itol.embl.de/). Color code represents different UGT groups. The novel UGT cluster of *Capsicum* is highlighted in pink. OG = outgroup.

## Discussion

Here we provide the first comprehensive analysis of the genetic architecture of pepper metabolism. To achieve this, we utilized two distinct populations. First, a natural variation-based GWAS panel of Balkan pepper which is widely used in commercial European breeding programs (Nankar et al., 2020). Secondly, we complemented the GWAS with a metabolite QTL analysis on a biparental BIL population (BC_2_S_4_) derived from *C. annuum* cv. Marconi giallo and *C. chinense* cv. CGN22793 (Fig. S3).

Our studies revealed that *Capsicum* harbors a diverse metabolic landscape with metabolite variation spanning the range of absent (below the threshold of detection) to 150-fold more abundant with this span being similar to that recorded in previous more targeted metabolite-based research (Alseekh et al., 2015; Brog et al., 2019). Results from the mGWAS were not affected by confounding effects of population structure (Fig. 1), and additionally, a subset of the identified loci from the mGWAS and mQTL analyses co-map either to loci previously associated with pungency (Fig. S7) or with earlier targeted metabolite analyses that identified transgressive segregation of metabolite production (Fig. S10, Data S8). These findings suggest that detailed characterization of these loci could reveal the molecular basis of both fruit pungency and metabolite heterosis. Indeed, in the case of the former, we were able to provide considerable insight in our study by being able to narrow down the previously mapped region QTL for capsaicinoids and fruit pungency on chromosome 2 (Tripodi et al., 2021; Cao et al., 2022) to a 28 gene interval including the gene *CA02g19260* encoding *AT3/PUN1* (Qin et al., 2014) and MYB transcription factor (*CA02g20190*).

Beyond the above example, the combined panels uncovered a further 93 candidate gene with potential involvement in pepper metabolism. This included a major cluster of 23 1-aminocyclopropane-1-carboxylate oxidases (ACO) involved in ethylene signaling, which are known regulators of stress responses and developmental processes (Houben and Van de Poel, 2019). To highlight some additional genes: *CA06g22860* is predicted to be a capsanthin/capsorbin synthase and could be identified on chromosome 6 using the BIL population. Further, a phytoene synthase (*CA04g04080*) on chromosome 4 which is associated with capsanthin and capsorbin derivatives, and a carotenoid cleavage dioxygenase (*CA11g03240*) on chromosome 11, could be identified (Data S9). Finally, seven QTL involved in the capsianosides pathway were identified (Fig. 3). Two terpenoid synthases (TPS) on chromosomes 3 and 11 are responsible for mono-/tri-/sesqui-/di- and tetraterpenes (Christianson, 2017), while a geranylgeranyl diphosphate reductase on chromosome three is responsible for synthesizing photosynthesis related isoprenoids while consuming GGPP (Ruiz-Sola et al., 2015).

In addition to the above-mentioned QTL, 11 UGTs form a significant hotspot region on chromosome 9 that is primarily responsible for the chemical diversity of capsianosides. Amongst the metabolites found to associate with this hotspot are acyclic diterpenoid glycosides. These compounds possess anticancer and defence-related potential and are unique to the genus *Capsicum* (Izumitani et al., 1990; Chilczuk et al., 2020; Bacon et al., 2017; Hashimoto et al., 1997). This major QTL is specific to pepper fruit pericarp (Fig. 2) and is additionally evidenced by mapping PC1 of capsianoside derivatives in all three panels (Figs. 4, S9, S10).

Because it is unknown whether differential glycosylation of capsianosides affects fruit morphology or taste, it is possible that breeders selected primarily for disease resistance (as evidenced by the presence of 74 disease resistance-related genes in close proximity), and that the UGT cluster was hitchhiking to support hypersensitive resistance against pathogens.

Wahyuni *et al*., (2013) previously demonstrated a clear separation among 32 accessions from four Annuum clade members based on differential glycosylation levels of capsianosides (Wahyuni et al., 2013), which is partially reflected in PC1 (Fig. S9). In a subsequent study, they used 113 F_2_ plants to map a small subset of metabolic QTL, including a hotspot region containing hundreds of QTL o which some are associated with capsianoside V, IX, VII, X-2, a capsicoside, and several flavonoids (Wahyuni et al., 2014). Here, we report of a QTL with a mere 25 genes which is primarily responsible for differential glycosylation of capsianosides. Although well-correlated compounds such as oxylipins and capsicosides (Fig. S19, in total 267 mass features) also mapped to this region as well. The reliance of these classes on the same chromosomal interval may be due to metabolic network compensation (Wahyuni et al., 2014). Sterols are derived from IPP and oxylipins are derived from fatty acids that share acetyl-CoA as a precursor with the MVA pathway for IPP biosynthesis (Gabbs et al., 2015; Toll, 2014). As such, their abundance is likely to be influenced by capsianoside abundance that said the mechanism by which flux to these three compound classes are linked remains elusive. Intriguingly, abrogation of the nicotianoside pathway in *Nicotiana* results in non-specific hydroxylation, which leads to autotoxicity via inhibition of the sphingolipid pathway (Heiling et al., 2016; Li et al., 2021). Therefore, we believe that glycosylation solves the autotoxicity problem and that digestion by herbivores promotes hydroxylations (Li et al., 2021). It appears that the intermediate compounds from 17-hydroxygernayllinalool to capsianosides, known as lyciumosides (Terauchi et al., 1995), have mapped to the same hotspot region on chromosome 9 as capsianosides in this study. These intermediate structures have not been described in *Capsicum* species hitherto.

However, pleiotropy analysis clearly demonstrated that a subset of metabolic features from both the GWAS and the biparental panel mapped to a variety of small effect QTL. This finding reinforces the power of natural variation and BILs while highlighting a major limitation of segregating F_2_ populations in analyzing only a limited genetic repertoire at low resolution.

Also fascinating were the implications from Tajima’s D analyses (Fig. 5a), which indicated a positive selective sweep in *C. annuum* in the region of this hotspot with a higher variation observed than expected. Because of a recent population contraction, *C. annuum* cultivars have undergone purifying selection which enhanced variation, and ultimately led to a higher resistance by means of a more diverse capsianoside landscape.

UGTs play an important role in the formation of fruit quality, pigmentation, and resistance to many antimicrobials and diseases (Alseekh et al., 2020). The UGTs studied here could be characterized as potential di-/triglycoside forming GTs and displayed similarity to group O family UGTs (Figs. 7, S15, S16). Duplication and subsequent diversification could be one explanation for the two clusters based on amino acid sequence similarity. Finally, molecular validation procedures confirmed the role of the *Capsicum* UGT cluster in the decoration of monomeric capsianosides.

In conclusion, our finding suggests that *Capsicum annuum* accessions from the Balkans have been divided into three genetically homogeneous groups resulting from morphological selection during domestication and/or through historical trading routes. Metabolic profiling of 162 pepper accessions, as well as 100-odd BILs grown under two conditions revealed a wide range of metabolic variations some of which correlated with plant morphometric descriptors (Fig. S19). Moreover, a combination of GWAS and QTL mapping revealed hundreds of QTL regulating specialized secondary metabolites; with more than 37 QTL discovered in both panels, and in total yielding 93 candidate genes. These studies highlight the utility of both the GWAS collection and the novel BIL population that we introduce here for understanding fundamental and applied aspects of pepper biology. Perhaps most important amongst these is a hotspot region on chromosome 9 that regulates monomeric capsianoside glycosylation. We were able to resolve the underlying genetics to a narrow region containing a cluster of eleven UGTs. The identification of this cluster adds to the burgeoning list of clusters in plant secondary metabolism (Zhan et al., 2022; Nützmann et al., 2016). Its importance is underlined by the evidence of the selection of these genes during pepper domestication as well as the anticancer potential of capsianosides (Chilczuk et al., 2020). As such, we believe that we have identified an important tool that will enable breeders to breed highly resistant and nutritionally enhanced biofortified peppers. This study will additionally provide a useful blueprint for the identification of the genetic architecture of nutraceuticals in all our crops.

## Materials and Methods

### Plant materials, sequencing, and SNP identification

A collection of 162 *Capsicum annuum* accessions from the Balkans (Albania, Bulgaria, Greece, Macedonia, Romania, and Serbia) was grown in a field plot trial in 2017 and 2018 in a randomized complete block design with three replications at the ‘Maritsa’ Vegetable Crop Research Institute, Plovdiv, Bulgaria. Seeds were sown at the end of March in an unheated greenhouse and transplanted in the middle of May in a two-rowed planting scheme. Plant protection, irrigation, microclimate, and fertilization were the same for all accessions. Phenotypic traits and fruit compositional quality were examined as previously described by Nankar *et al*., (2020). Genotyping was performed following the genotyping-by-sequencing (GBS) approach which resulted in 1,149,249 SNPs, by applying standard filtering with MAF 5 % we end up with 77,401 SNPs across the population which after used to conduct the GWAS analysis (Fig. 1a).

An interspecific backcross inbred line (BIL) population of 117 lines was generated and used in this study. The BIL population was developed from a cross between *C. annuum* cv. Marconi giallo and *C. chinense* cv. CGN22793 through two generations of backcrossing using *C. annuum* cv. Marconi giallo as the recurrent parent followed by four generations of selfing. The 117 BILs and their two founder lines were planted in two environments (glasshouse and polytunnel) at the Research Centre for Vegetable and Ornamental crops (Pontecagnano, SA, Italy, Latitude: 40°38′59.8“N; Longitude: 14°53′31.6”E). The 117 lines were arranged in a randomized complete block design with three replications and a two-rowed planting scheme. BILs were grown from May to October 2019 under the same conditions. DNA was extracted from all the lines and subjected to GBS. In total 10,914 polymorphic SNPs were identified and used to perform the QTL mapping. SNP discovery and annotation were performed as previously described (Taranto et al., 2016). The genomic composition of the BIL population was analyzed using the software CSSL Finder (http://mapdisto.free.fr/CSSLFinder/) and GGT2.0 (Van Berloo, 2008).

### LC-MS metabolite profiling

Mature fruits were harvested and pericarp tissue was either lyophilized for the association panel or freshly processed by grinding under frozen conditions (28 Hz, 30 sec). Fresh fruit materials (120 mg) and freeze-dried materials (30 mg) were extracted by MTBE method as previously described (Giavalisco et al., 2011; Salem et al., 2016) spiked with 12.5 µg/ml of isovitexin. The semi-polar phase (120 µl and 240 µl) were taken to dryness using a vacuum centrifuge. Analysis of semi-polar metabolites was conducted on a UPLC-LC-MS machine as described previously (Alseekh et al., 2015). In brief, the UPLC system was equipped with an HSS T3 C18 reverse-phase column (100 × 2.1 mm internal diameter, 1.8 μm particle size; Waters) that was operated at a temperature of 40°C. The mobile phases consisted of 0.1 % formic acid in water (solvent A) and 0.1 % formic acid in acetonitrile (solvent B). The flow rate of the mobile phase was 400 μL/min, and 3 μL of the sample was loaded per injection. The UPLC instrument was connected to an Exactive Orbitrap focus (Thermo Fisher Scientific) via a heated electrospray source (Thermo Fisher Scientific). The spectra were recorded using full-scan in negative ion-detection mode, covering a mass range from m/z 100 to 1,500. The resolution was set to 70,000 and the maximum scan time was set to 250 ms. The sheath gas was set to a value of 60 while the auxiliary gas was set to 35. The transfer capillary temperature was set to 150 °C while the heater temperature was adjusted to 300 °C. The spray voltage was fixed at 3 kV, with a capillary voltage and a skimmer voltage of 25 V and 15 V, respectively. MS spectra were recorded from 0 to 19 minutes of the UPLC gradient. Processing of chromatograms, peak detection, and integration was performed using RefinerMS (version 5.3; GeneData). Metabolite identification and annotation were performed using standard compounds, tandem MS (MS/MS) fragmentation, and metabolomics databases. When using the in-house reference compound library we allowed for a 10 ppm mass error and a dynamic retention-time shift of 0.1. Metabolite data is reported following recently updated standards for metabolite reporting (Alseekh et al., 2021).

### Population structure analyses

To analyze the population structure underlying the GWAS population the software Structures 2.3.4. (Pritchard et al., 2000) was utilized following the admixture model with correlated alleles with K 2 – 8. Six independent runs with 500 burn-in generations and 5000 Markov Chain Monte Carlo generations were used for estimating K. The optimal K value was determined by ΔK (Evanno et al., 2005) estimated with PopHelper (Francis, 2017). Principal component analysis, dendrogram, correlation matrix, and heatmaps were either designed by Genedata Expressionist® Analyst software or MetaboAnalyst 5.0 (Pang et al., 2021).

### Genome-wide association analyses

The R package rMVP (Yin et al., 2021) was used employing a genomic relationship matrix, efficient mixed-model association, α-threshold of 0.05, and mixed linear model. Manhattan plots were employed by the package qqman (Turner, 2018) with a genome-wide threshold calculated with Bonferroni correction (-log10(0.05/77401)) and a suggestive threshold approximating false-discovery rate (-log10(1/77401)) mapped against the reference genome sequence *C. annuum* cv. CM334 version 1.6 from PLAZA 4.0 (Van Bel et al., 2018).

### QTL mapping

R/qtl (Broman et al., 2003) was used to construct a linkage map using a permutation rate of 1000 and alpha = 0.01, and a logarithm of odds (LOD) threshold of 15 was defined for the relative intensities of 2551 mass features.

### Statistical analyses

Broad-sense heritability was calculated using the following equation by treating independent accessions and BILs as a random effect and the biological replication as a replication effect: H^2^ = var_(G)_/(var_(E)_+var_(G)_), where var_(G)_ and var_(E)_ is the variance derived from genetic and environmental effects, respectively. The variation across accessions and BILs was determined by the first and third quartiles. Tajima’s D in 1 Mb non-overlapping bins was calculated using VCFtools (Danecek et al., 2011). Linkage disequilibrium was estimated by squared allele-frequency correlation (r^2^) according to the R package LDheatmap (Shin et al., 2006) in a ± 50 kb window. Orthologous genes and orthogroups were discovered by OrthoFinder (Emms and Kelly, 2018, 2015). Protein sequences were obtained from genome databases: PLAZA 4.0 (https://bioinformatics.psb.ugent.be/plaza), peppergenome (peppergenome.snu.ac.kr), SolGenomics (solgenomics.net) and TAIR (arabidopsis.org). Pleiotropy analysis was conducted by the circos representation Fuji plot (Kanai et al., 2018; Krzywinski et al., 2009) using 150 and 136 specialized metabolic features and 1,780 and 1,177 SNPs of association and biparental panel based on their putative annotation and H^2^ higher 95 %. Locus-IDs were assigned in a ± 50 kb window.

### Phylogenetic analysis

Sequence alignment of BIL and GWAS individuals was performed by MAFFT version 7 (https://mafft.cbrc.jp/alignment/server/), and its visualization by iTOL (https://itol.embl.de/). Sequences for the multiple sequence alignment of the UGTs were obtained from Knoch *et al*., 2018; Wilson and Tian, 2019; Heiling *et al*., 2021; Dudley *et al*., 2022(Knoch et al., 2018; Wilson and Tian, 2019; Heiling et al., 2021; Dudley et al., 2022). Used abbreviations: *UGT1* (*scaffold551.50*), *UGT2* (*scaffold551.53*), *UGT3* (*scaffold551.54*), *UGT4* (*scaffold551.55*), *UGT5* (*scaffold551.57*), *UGT6* (*scaffold551.58*), *UGT7* (*scaffold551.60*), *UGT8* (*scaffold551.61*), *UGT9* (*scaffold551.62*), *UGT10* (*scaffold551.63*) and *UGT11* (*scaffold551.64*).

### Transient overexpression

The total RNA from *C. annuum* cv. Marconi giallo and *C. chinense* cv. CGN22793 was isolated using a NucleoSpin® RNA plant kit (Macherey-Nagel) according to the manufacturer’s instructions. First-strand cDNA was synthesized using 1.5 µg RNA and Prime Script™ RT reagent Kit with gDNA eraser (Takara) according to the manufacturer’s instructions. Full-length cDNA of fruit pericarp samples of the founder lines was amplified by using 250 ng (primers see Data S15). The entry clone was obtained through recombination of the PCR product with pDONR207 (Invitrogen). By LR recombination error-free clones were introduced into pK7FWG2 (Karimi et al., 2002). Transformation of leaves of *C. annuum* cv. CM334, *N. benthamiana,* and *A. thaliana* were performed following (Zhang et al., 2020b), and *Agrobacterium tumefaciens* (AGL1) containing vector pBin61-p19 infiltrated with an OD_600_ 0.5. DM6000B/SP5 confocal laser scanning microscope (Leica Microsystems, Wetzlar, Germany) was used for verification of expression. Metabolic shifts were analyzed after 1 and 3 days using 50 mg (*C. annuum*, *N. benthamiana*) and 20 mg (*A. thaliana*), respectively, of agro-injected leaves and subjected to LC-MS analysis as described above using chloroform:methanol:water extraction (Bligh and Dyer, 1959).

## Data availability statement

The data are provided in the supplemental. More information is available upon request.

## Author Contributions

S.A. and A.R.F. conceptualized the experiment. J.N., A.R.F. and S.A. wrote the manuscript with input from all authors. J.N. and P.T. performed experiments, and J.N. and S.A. performed metabolite profiling. J.N. performed data analysis with the support of R.W., A.K., and M.B., and Y.T., and A.B. contributed analysis tools. V.Tr., A.N.N., V.S., G.P., V.To., T.G., and D.K., and P.T developed the GWAS panel and the BIL population, conducted field trials and provided plant materials.

## Acknowledgments

We wish to acknowledge the G2P-SOL project, VT, AN, VS, GP, TG, DK, ARF, and SA acknowledge the European Union’s Horizon 2020 research and innovation program, project PlantaSYST (SGA-CSA No. 739582 under FPA No. 664620) and project BG05M2OP001-1.003-001-C01. The European Union Horizon 2020 research and innovation program under Grant Agreement No. 677379 and the European Regional Development Fund through the Bulgarian “Science and Education for Smart Growth” Operational Program.

